# Unique transcriptomic landscapes identified in idiopathic spontaneous and infection related preterm births compared to normal term births

**DOI:** 10.1101/386185

**Authors:** Heather M Brockway, Suhas G Kallapur, Irina A Buhimschi, Catalin S Buhimschi, William E Ackerman, Louis J Muglia, Helen N Jones

**Author notes:** Joint senior authors (Louis J Muglia, Helen N Jones).

## Abstract

Preterm birth (PTB) is leading contributor to infant death in the United States and globally, yet the underlying mechanistic causes are not well understood. Histopathological studies of preterm birth suggest advanced villous maturity may have a role in idiopathic spontaneous preterm birth (isPTB). To better understand pathological and molecular basis of isPTB, we compared placental villous transcriptomes from carefully phenotyped cohorts of PTB due to infection or isPTB between 28-36 weeks gestation and healthy term placentasu. Transcriptomic analyses revealed a unique expression signature for isPTB distinct from the age-matched controls that were delivered prematurely due to infection. This signature included the upregulation of three IGF binding proteins (*IGFBP1, IGFBP2,* and *IGFBP6*), supporting a role for aberrant IGF signaling in isPTB. However, within the isPTB expression signature, we detected secondary signature of inflammatory markers including *TNC*, *C3*, *CFH,* and *C1R*, which have been associated with placental maturity. In contrast, the expression signature of the gestational age-matched infected samples included upregulation of proliferative genes along with cell cycling and mitosis pathways. Together, these data suggest an isPTB molecular signature of placental hypermaturity, likely contributing to the premature activation of inflammatory pathways associated with birth and providing a molecular basis for idiopathic spontaneous birth.

## Introduction

Every year, approximately 15 million infants are born before 37 completed weeks of gestation and of those, 1.1 million die from complications resulting from their premature birth. In 2018, the incidence of prematurity in the United States was 10.4%[1]and worldwide incidences reached approaching 20% in some regions[2]. There are two primary clinical classifications: spontaneous preterm birth (sPTB) and iatrogenic preterm birth as a result of fetal or maternal complications. Advances in the study of sPTB have allowed the distinction of births due to infection or truly idiopathic spontaneous preterm birth (isPTB). Yet, these broad classifications do not adequately describe the heterogeneous nature of preterm birth nor have they allowed for a systematic understanding of the underlying molecular mechanisms driving the observed heterogeneity. This has led to a lack of diagnostics and effective therapeutic interventions in the clinical setting.

While risk factors such as diet, environmental exposures, and a family history have been identified, the underlying molecular mechanisms of sPTB remain unclear[3]. Furthermore, the molecular differences between infection and isPTB are not well understood. However, one key to understanding the heterogeneity associated with preterm birth may be the placenta, a transient two-sided organ, contributing to the maternal/fetal interface, and along with its proper development and function is essential to a successful pregnancy outcome[4]. Placentation is the result of a highly complex web of molecular interactions originating from both the mother and the fetus. We do not fully understand these processes in healthy, term pregnancies let alone their impact on adverse pregnancy outcomes[4–6]. However, recent studies utilizing placental pathology have started to provide insights into the morphological differences in preterm birth.

Morgan et al., conducted histological studies of the villous trophoblast in isPTB and infection (<37 weeks GA) and identified four distinct morphological subclasses between isPTB and infection. Within the isPTB samples, one subclass, comprising 59% of the samples, had morphological hallmarks indicative of term placenta and was classified as having advanced villous maturation (AVM)[7]. The second subclass had none of these hallmarks and made up the remaining 41% of the samples examined[7–9]. These data suggested over half of isPTB samples were associated with hypermature placentas. These authors also examined a set of infection samples and observed that in contrast to the isPTB, 19% had hallmarks of AVM in addition to infection while the remaining 81% had only infection. A second study by Nijman et al., also examined placental villous histology in spontaneous preterm births between 24 weeks and 36.6 weeks gestation using morphological hallmarks associated with maternal villous malperfusion (MVM) of which AVM is a specific subtype[7,10]. Again, spontaneous preterm samples were associated with four different morphologies: only hallmarks of MVM, only hallmarks of infection, hallmarks of both MVM and infection, and with some samples having no distinct morphological hallmarks at all, thus confirming the findings of the Morgan et al study. Interestingly, these morphologies tracked with gestational age; with samples between 24-28 weeks having predominately infection hallmarks alone or MVM and infection hallmarks, while those samples between 28-36.6 weeks had predominately either MVM hallmarks or none at all[10]. Together, these two studies suggest at least four subclasses of sPTB exist based on histological observations of the placental villous tissue and these subclasses are likely contributing to the heterogeneity confounding identification the underlying etiology of isPTB.

We know from molecular studies of normal term pregnancies, there is coordinated development and maturation between the fetus and the placenta throughout gestation with placental villous remodeling occurring between 20-40 weeks[5]. This remodeling leads to the increase in terminal villi density which reflects an enlarged placental surface area, allowing for increasing nutrient transport to accommodate the growing fetus. Given the morphological evidence for hypermaturity of the placental villous tissue, we hypothesize there is a developmental disconnect occurring in a subset of isPTB pregnancies where the molecular mechanisms associated with normal maturation are aberrant resulting in the hypermaturity observed in the Morgan and Nijman studies, resulting in the premature initiation of parturition resulting in preterm birth. We further hypothesize placentas with infection lesions lack this developmental disconnect as the infection is likely inducing activation of inflammatory pathways associated with parturition initiation, resulting in early delivery. To test these hypotheses and to identify placental molecular mechanisms of preterm subtypes, we obtained histologically phenotyped placental villous samples from isPTB and acute histologic chorioamnionitis(AHC) births between 29 and 36 weeks and normal term births between 38 and 42 weeks, generated transcriptomes and through comparative computational analyses, identified distinct molecular signatures for isPTB and AHC births.

## Results

### Study Cohort

Maternal and fetal characteristics for the three different pregnancy outcomes included in the transcriptomic analyses are presented in Table 1. Significant differences were observed in gestational age and fetal weights between AHC and isPTB samples compared to the term samples (P<0.05). Among the isPTB samples, two neonates were small for gestational age (SGA) with a fetal weight less than the 10^th^ percentile with the remaining neonates above the 50^th^ percentile for birth weight. All AHC and term births for which there were fetal weights available were appropriate for gestational age. We included males and females in each sample set and adjusted the linear models for fetal sex in addition to birth outcome. It is important to note that in this study, we have mixed ethnic background within each of the sample sets.

**Table 1:** Clinical characteristics of the placental villous samples included in the transcriptome analyses

| Characteristics | Acute<br>Histologic<br>Chorioamnionitis<br>Births<br>(AHC) | Idiopathic<br>Spontaneous<br>Preterm Births<br>(isPTB) | Term Births | P values |
| --- | --- | --- | --- | --- |
| Number of samples | 9 | 12 | 11 |  |
| Maternal Age | 33(25-40) | 28(18-39) | 30(19-37) | NS<sup>1</sup> |
| Gestational Age | 31(29-34)* | 33(30-36)* | 39(38-42) | <0.0001<sup>1</sup> |
| Fetal sex (% female) | 4(44%) | 6(50%) | 4(36%) | NS<sup>3</sup> |
| Fetal weight (grams) | 1820(1360-2300)* | 2062(1450-2722)* | 3795(3650-4527) | <0.05<sup>2</sup> |
| Birth weight percentile | 60(20-80) | 45(3-80)* | 80(60-99) | <0.05<sup>2</sup> |
| SGA % | 0 | 16.6% | 0 |  |
| <u>Delivery type</u> |  |  |  |  |
| Cesarean (%) | 4(44%) | 5(42%) | 6(54%) | NS<sup>3</sup> |
| <u>Infection Status</u> |  |  |  |  |
| (% Positive) | 9(100%)* | 0 | 1(7%) | <0.0001<sup>3</sup> |

Data shown as median with range or total number with percent
<sup>1</sup> ANOVA with Tukey's correction for multiple comparisons
<sup>2</sup> Kruskal Wallis ANOVA with Dunn's correction for multiple comparisons
<sup>3</sup> Chi Square Analyses NS=Not significant
\*indicates the group significantly different from Term births.

### Identification of significant differentially expressed genes

Due to the origin of the samples and inclusion of a pre-existing dataset, we performed multiple assessments to identify potential batch effects which would confound downstream analyses and determined there were no significant batch effects identified. Thus, we did not remove or control for them in subsequent models or tests for differential gene expression. The data matrix was filtered and normalized within edgeR (Emperical Analyses of Digital Gene Expression in R)[11], leaving us with a total of 13,929 genes in the data matrix for analysis.

To account for the type of birth and fetal sex differences, we utilized the generalized linear modeling function (glm) within edgeR. Once the model was built, we conducted differential gene expression testing using three pairwise comparisons: isPTB compared to term births (TB), AHC compared to isPTB, and AHC compared to TB. Genes were considered significant with an adjusted P-value of <0.05 and absolute log2 fold-change >1.0 (Figure 1A). In the isPTB vs TB comparison, we identified a total of 188 significant differentially expressed genes, with 183 upregulated and only 5 downregulated. The AHC vs isPTB comparison, representing the samples of the same gestational age, yielded a total of 347 significant differentially expressed genes, with 96 upregulated and 251 downregulated. In the final comparison, AHC vs TB, we identified 187 significant differentially expressed genes with 137 upregulated and only 50 downregulated (Figure 1A). To identify transcriptomic signatures due to isPTB or AHC pathophysiology, we needed to categorize potential candidate genes as those with an isPTB or AHC expression pattern. We intersected the genes, both upregulated and downregulated, from each of the differential gene expression comparisons (Figure 1B). We were able to parse the genes into AHC and isPTB categories as well as some that did not fit either category but appeared to be gestationally age related.

**Figure 1:**
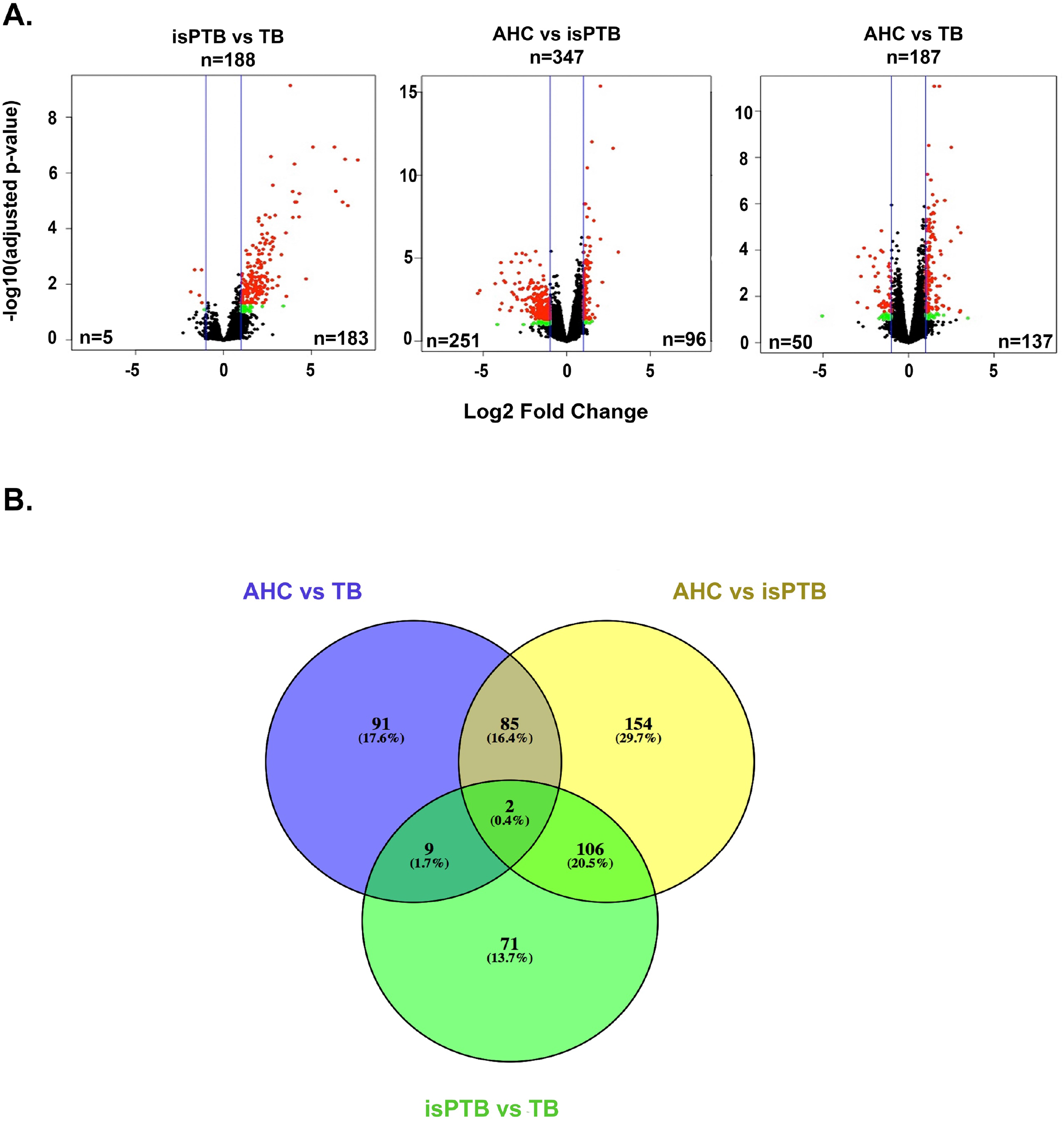
A comparative approach to identifying molecular signatures of isPTB and AHC. **A.** Differentially expressed genes were identified using pairwise comparisons *edgeR*. Red dots represent significant genes that have an absolute log2 fold-change of 1.0 and Benjamini Hochberg adjusted P-value of <0.5. Green dots represent genes with an absolute log2 fold-change of 1.0 and Benjamini Hochberg adjusted P-value of <0.1. Blue lines represent log2 fold-change of 1.0. **B.** The Venn diagram represents the intersection of significant genes from Panel A which was utilized to further refine the classification into molecular signatures for isPTB and AHC.

### The transcriptomic signature of isPTB is distinct from AHC and normal term samples

We furthered refined our analyses by assessing the expression pattern of each candidate genes in each group across all three differential expression datasets (Figure 2A and B). To be categorized as an isPTB candidate, a gene needed to be upregulated/downregulated in the isPTB compared to AHC or TB, and ideally demonstrating a non-significant difference in the AHC vs TB comparison (Log2 fold-change <1). We identified total of 170 genes that generally fit this expression pattern, all upregulated in their respective isPTB comparisons (Supplemental Table S2). 102 of these genes had an expression pattern where the fold-changes were greater in the isPTB vs TB comparison than in the isPTB vs AHC comparison. Not all genes AHC vs TB comparison fit our criteria for a non-significant or no difference. Several genes do have a significant difference in that comparison although it was to a lesser extent compared to the isPTB comparisons, indicating an underlying basal expression level in these genes during gestation. Upregulation in the AHC vs TB comparison indicates high expression earlier in gestation that declines with advancing gestational age (Figure 2A). The remaining 68 genes had a slightly different expression pattern, where the fold-change in expression between isPTB and AHC was greater than fold-change in expression between isPTB and TB and with essentially no significant difference in the TB vs AHC comparison, suggesting these genes belong to a specific isPTB molecular signature (Figure 2B). Given the isPTB and AHC comparison represents a comparison between overlapping gestational ages, this expression pattern suggests a hypermaturity molecular signature. We also observed that in this group of genes, that upregulation in the TB vs AHC comparison suggestion expression earlier in gestation that increases with advancing gestational age (Figure 2B).

**Figure 2:**
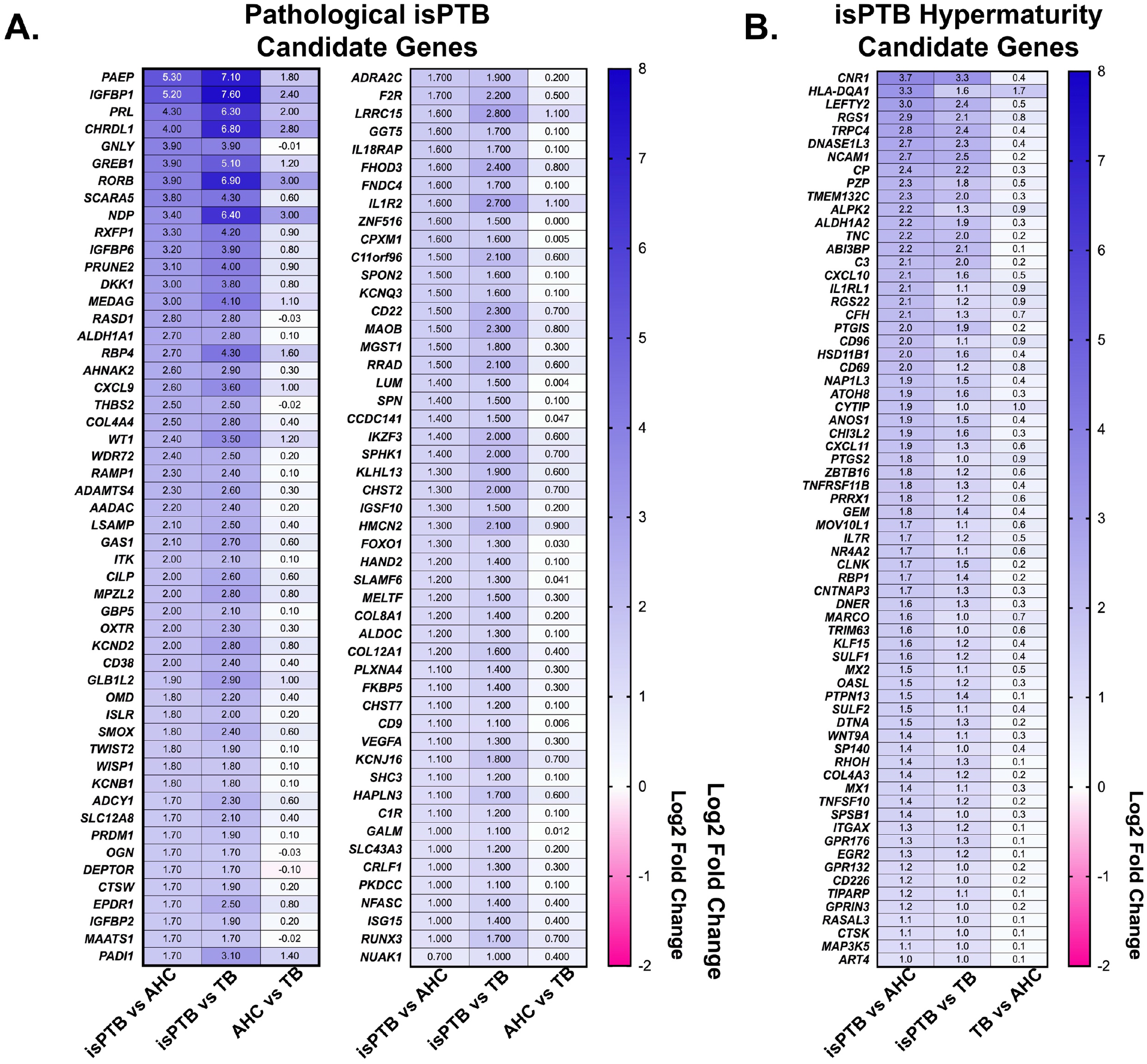
Identification of isPTB specific transcriptomic signature. **A.** Pathological isPTB candidate genes were identified by assessing the differential expression across all three pairwise comparisons. isPTB candidate genes had significant upregulated expression in isPTB samples compared to TB and AHC, with the either no change in expression pattern between AHC and TB or upregulation in AHC, although to a lesser extent than observed in the isPTB samples. This slight upregulation in AHC samples indicates a baseline expression is higher in samples of earlier gestational age and decreases over term. **B.** isPTB hypermaturity genes were also identified in the same manner as those in Panel A with the reversal of expression assessment between AHC and TB. In this analysis, we observed that the expression of these genes increases over gestation with expression being increased in TB samples compared to AHC. More importantly, when comparing isPTB to AHC which overlap in gestational age, there is a significant upregulation in isPTB samples which is suggestive of hypermaturity. Genes are arranged in order of Log2 fold change in the isPTB vs AHC comparison. Values = Log2 fold change

### IGFBP1 and DKK1 are upregulated in syncytiotrophoblasts in isPTB

We validated upregulation of two isPTB candidate genes, *IGFBP1* and *DKK1* using immunohistochemistry (IHC) on TB and isPTB placental samples. Both proteins localize to the syncytiotrophoblast in TB samples with a marked increase in expression in the isPTB samples (Figure 3). The reduced expression in the term tissues confirms the observations made in the isPTB expression data that there is basal expression of these genes during gestation.

**Figure 3:**
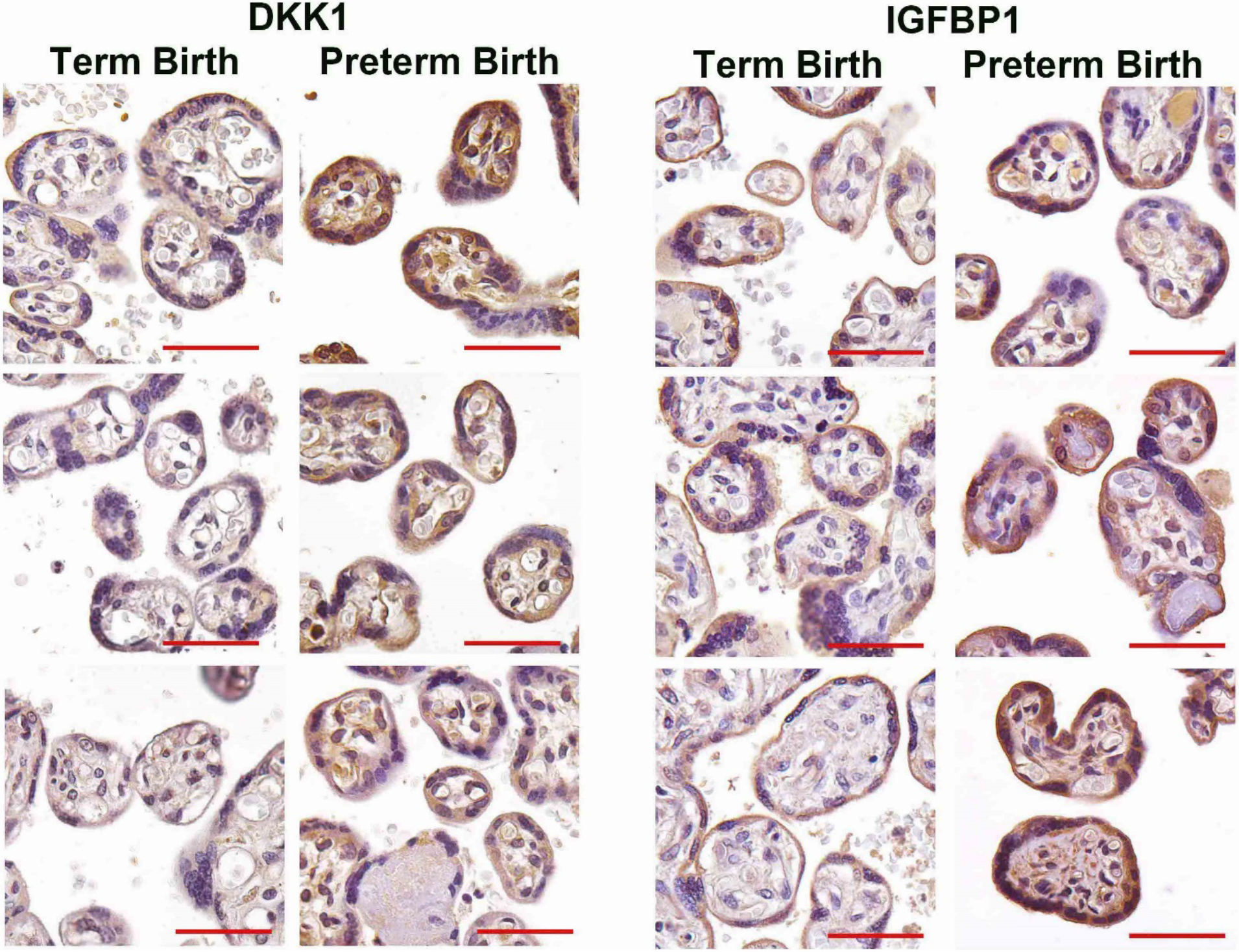
isPTB placental tissue samples demonstrated increased DKK1 and IGFBP1 expression. DKK1 and IGFBP localization the syncytiotrophoblast in the control term births with increased expression in isPTB. Images are taken at 40x magnification and scale bar = 50um

### The AHC transcriptomic signature does not overlap with the isPTB signature

We conducted a similar categorization of AHC genes (Figure 4) where the expression in the AHC comparisons were upregulated or downregulated compared to isPTB and TB which were expected to show a no difference in expression. We identified 170 genes that do not overlap with the isPTB candidates, representing a distinct AHC transcriptomic signature (Supplemental Table S3). The AHC signature includes 137 upregulated genes and unlike the isPTB signature, 33 downregulated genes (Figure 3). Within the isPTB vs TB comparison, there are no genes that are differentially expressed, indicating a similar expression pattern within these respective birth types.

**Figure 4:**
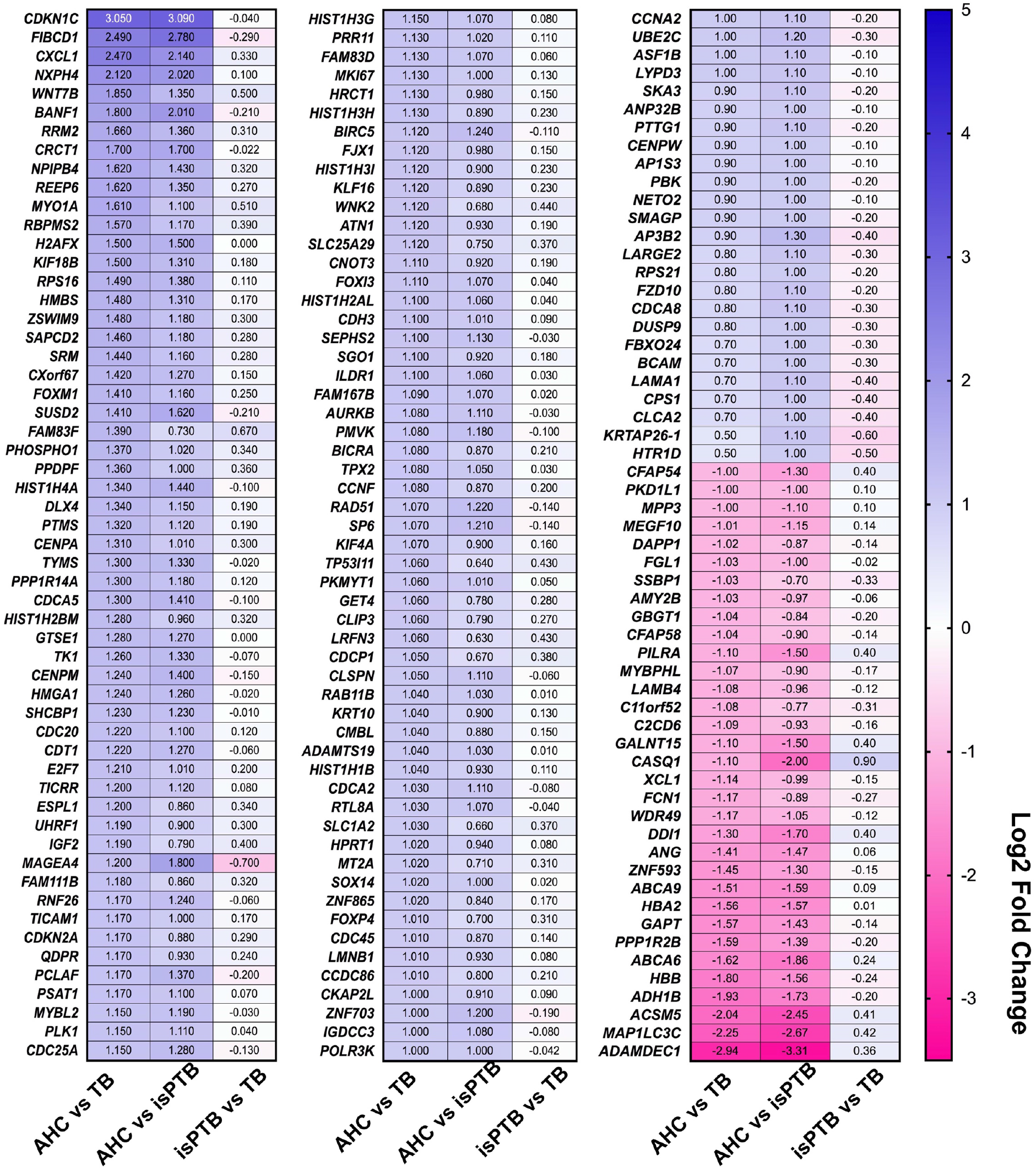
Identification of an AHC transcriptomic signature. AHC candidate genes were identified by assessing the expression pattern across all three pairwise comparisons. In this instance, we observed greater differential expression, both upregulated and downregulated, in the AHC samples compared to isPTB or TB with either no difference or non-significant differences in isPTB vs TB comparisons. Genes are arranged in order of Log2 fold change in the AHC vs TB comparison. Values = Log2 fold change

### isPTB candidate genes represent upregulated growth and inflammation pathways

We were able to identify molecular pathways of interest by analyzing our isPTB candidate genes lists through statistical overrepresentation. Our analysis of the isPTB candidate genes returned four significant pathways (Table 2). Of these pathways, two are directly associated with specific signaling pathways: the regulation of IGF uptake and transport by IGFBPs and cytokine signaling with the remaining pathways being more generalized to the immune system and signal transduction.

**Table 2:**
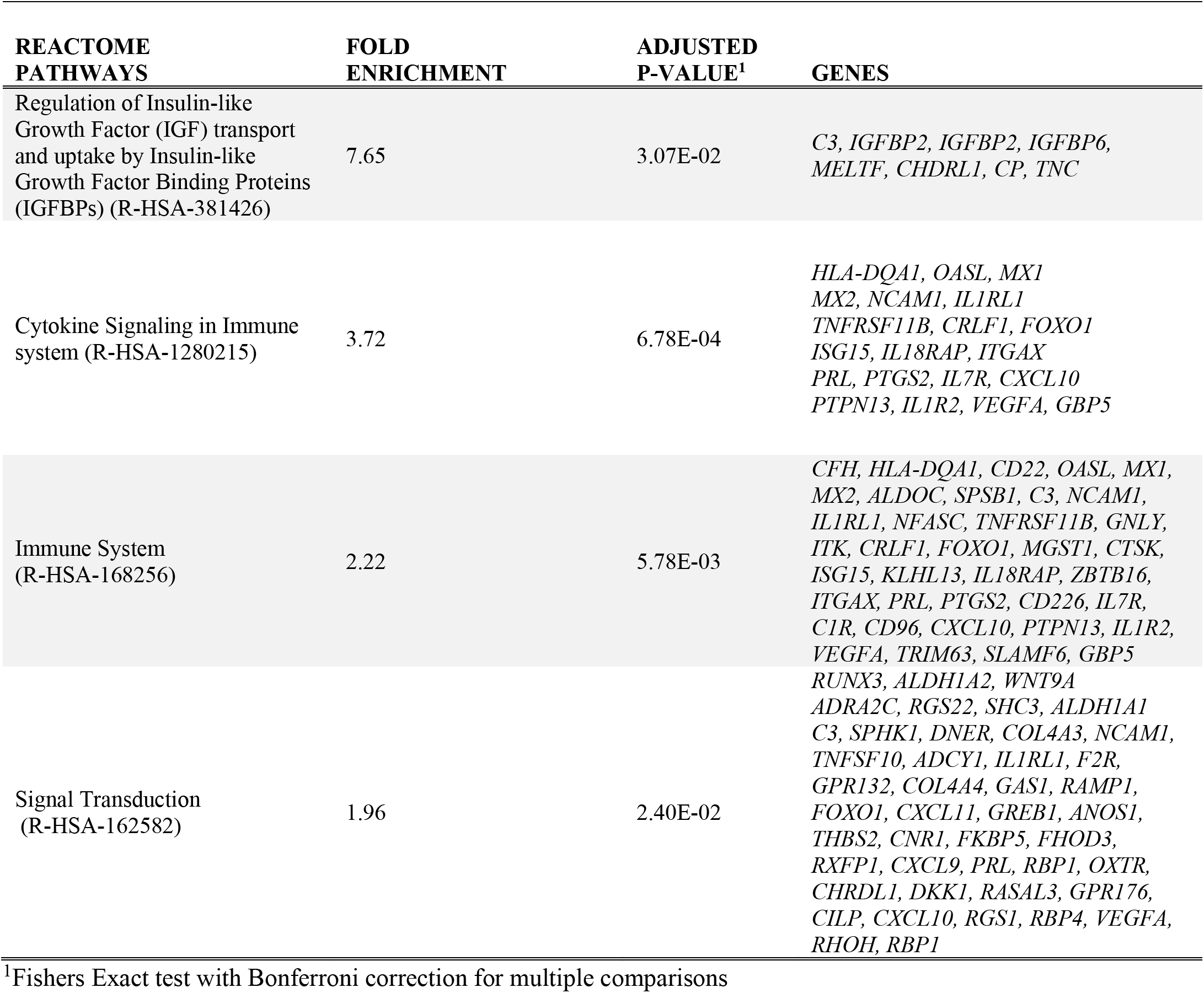
Reactome pathway enrichment analyses for isPTB candidate genes

### AHC candidate genes indicate increased mitotic activity

We conducted pathway analysis of the AHC candidate genes using statistical over-representation and identified 22 significant pathways (Table 3). 20 pathways are associated with regulation of cell cycling and mitosis (Table 3). Many of the genes overlap with multiple pathways. The remaining two pathways are associated with general cellular senescence and more specifically, DNA damage/telomere stress induced senescence.

**Table 3:** Reactome pathway enrichment analyses for AHC candidate genes

| REACTOME<br>PATHWAYS | FOLD<br>ENRICHMENT | ADJUSTED<br>P VALUE<sup>1</sup> |
| --- | --- | --- |
| Polo-like kinase mediated events (R-HSA-156711) | 38 | 1.14E-03 |
| Cyclin A/B1/B2 associated events during G2/M transition (R-HSA-69273) | 25 | 7.53E-03 |
| Activation of E2F1 target genes at G1/S (R-HSA-539107) | 22 | 1.23E-02 |
| G1/S-Specific Transcription (R-HSA-69205) | 22 | 1.23E-02 |
| DNA Damage/Telomere Stress Induced Senescence (R-HSA-2559586) | 14 | 2.41E-03 |
| Deposition of new CENPA-containing nucleosomes at the centromere (R-HSA-606279) | 14 | 1.70E-02 |
| Nucleosome assembly (R-HSA-774815) | 14 | 1.70E-02 |
| Amplification of signal from unattached kinetochores via a MAD2 inhibitory signal (R-HSA-141444) | 11 | 2.88E-03 |
| Amplification of signal from the kinetochores (R-HSA-141424) | 11 | 2.88E-03 |
| Mitotic Spindle Checkpoint (R-HSA-69618) | 10 | 8.73E-04 |
| Resolution of Sister Chromatid Cohesion (R-HSA-2500257) | 9 | 2.27E-03 |
| Mitotic Anaphase (R-HSA-68882) | 9 | 2.89E-06 |
| Mitotic Metaphase and Anaphase (R-HSA-2555396) | 9 | 3.08E-06 |
| Mitotic G1-G1/S phases (R-HSA-453279) | 8 | 1.22E-03 |
| Cell Cycle Checkpoints (R-HSA-69620) | 8 | 4.25E-08 |
| Separation of Sister Chromatids (R-HSA-2467813) | 8 | 1.43E-04 |
| Cell Cycle, Mitotic (R-HSA-69278) | 8 | 5.55E-15 |
| RHO GTPases Activate Formins (R-HSA-5663220) | 7 | 4.10E-02 |
| Cell Cycle (R-HSA-1640170) | 7 | 1.94E-15 |
| Cellular Senescence (R-HSA-2559583) | 7 | 2.03E-02 |
| M Phase (R-HSA-68886) | 6 | 1.63E-05 |
| RHO GTPase Effectors (R-HSA-195258) | 5 | 8.22E-03 |

<sup>1</sup>Fishers Exact test with Bonferroni correction for multiple comparisons

## Discussion

Understanding the etiology of each of the subtypes of preterm birth previously identified by studies in placental histopathology [8–10] will allow for better clinical diagnosis and treatment. We conducted three pairwise comparisons between placentas from preterm births within a linear model accounting for fetal sex and gestational age, then isolated common and unique signatures to isPTB and AHC. We further refined the unique signatures into specific lists of significant candidate genes then examined the lists to identify potential molecular pathways of interest to the pathophysiology of isPTB.

Within the unique isPTB signature, we observed significant over-representation of genes in insulin like growth factor signaling pathway: *IGFBP1*, *IGFBP2*, *IGFBP6*, *C3, MELTF, CP, TNC*, and *CHDRL1*. The upregulation of the IGFBPs could have multiple paracrine and autocrine implications throughout the context of an isPTB pregnancy[12,13]. Secreted IGFBP1 and IGFBP2 have a high affinity for IGF1 while IGFBP6 preferentially binds IGF2, reducing their bioavailability to activate IGF signaling [14]. The aberrant increase in *IGFBP1, IGFBP2*, *and IGFBP6* in isPTB placentas may suggest a reduction in IGF signaling, however we do not see reduced fetal weight in the majority of our samples suggesting placental supply to maintain fetal growth via the mTOR pathway is not affected[15,16].

IGFBP2 and IGFBP6 have roles independent of IGF signaling. IGFBP2 has been associated with enhanced cell proliferation via extracellular interaction with EGFR and the activation of the STAT3 signaling pathways[17]. It can also translocate to the nucleus to act as a transcription factor promoting VEGF expression [20,21]. Interestingly, IGFBP2 has a non-canonical promoter comprised of four putative NFKB binding sites. NFKB has previously been implicated in the activation of pro-labor pathways through non-canonical signaling via activation of the STAT3 pathways[18]. It is possible that increased IGFBP2 is activating EFGR/STAT3 due to NFKB or other signaling resulting in increased placental maturation and the premature activation of pro-labor pathways and thus, isPTB.

Independent of its roles in IGF signaling, IGFBP6 can inactivate WNT signaling by blocking WNT binding to the FDZ and LRP receptors[19]. WNT signaling is essential to placental development through STB differentiation, and most likely, the suppression of NFKB signaling, limiting the initiation of pro-labor inflammatory pathways[20]. Increased IGFBP6 would abrogate WNT signaling and the associated biological processes. While the WNT/B-catenin and NFKB cross-talk has not been fully explored in villous tissue, the upregulation of non-canonical NFKB pathways has been associated with the activation of pro-labor inflammatory pathways in villous trophoblasts of normal placentas [18]. Thus, the upregulation of IGFBP6 could represent premature activation of pro-labor pathways. IGFBP6 also functions at the intercellular level through its binding of Ku-80. Ku-80 functions in non-homologous DNA repair and the interaction with IGFBP6 is associated with cell cycling[21]. Furthermore, Ku proteins including Ku-80 have been implicated in telomere maintenance including maintaining telomere length which is associated biological aging [21]. Once a critical shorten length is reached, senescence is activated along with a corresponding hyperactivity of NFKB, TNFα, IL6, and IFNγ all known to have roles in activation of pro-labor pathways[21,22]It has been suggested that trophoblast telomeres may be a biological clock by which the placenta “measures” time, but the evidence thus far has been conflicting [23–25] perhaps due to the lack of appropriate placental phenotyping. Elucidating the non-canonical roles of IGFBP2 and IGFBP6 in villous tissues could provide additional insights to the molecular mechanisms underpinning isPTB.

Our immunohistochemistry studies support the transcriptome data demonstrating both increased IGFBP1 and DKK-1 protein expression. DKK1 is a WNT signaling inhibitor and WNT signaling is essential to the development and maturation of the placenta potentially through the differentiation of trophoblasts[26]. Just as IGFBP6 could be modulating the WNT-NFKB crosstalk and signaling, so to could DKK1 as they both function to block WNT singling by binding to the receptor[27] and thus activate pro-labor inflammatory pathways.

*TNC, C3, CP, MELTF,* and *CHRDL1* upregulation maybe due to the activation of RAS signaling through either the IGF signaling pathway[12], or possibly through IGFBP2 activation of EGFR or cytokine activation of growth factor signaling. TNC, C3, CP represent inflammatory markers[28–31], while CHRDL1 is a marker of cellular aging and BMP4 agonist, likely signaling the arrest of BMP4 related cell proliferation [32,33]. *TNC* and *C3* are of particular interest due to their roles in physiological processes central to placental maturation. Furthermore, they have been extensively studied in the context of cancer progression and inflammatory processes, with diagnostics and potential therapeutics already in place or in development[28,34]. TNC modulates numerous biological processes such inflammation, branching morphogenesis, epithelial-mesenchymal transformations, mediation of cell signaling through interactions connecting ECM molecules to cell membrane receptors such as TRL4, FRZ, EGFR, and integrins[28]. The upregulation of TNC in our isPTB samples could be indicative of increased branching of the villi coupled with increased cell signaling through its interaction with cell membrane receptors. As we are only examining the endpoint of gestation, we are unable to determine if TNC expression is upregulated throughout gestation, leading to premature branching and thus, hypermaturation. Given the lack of study in the placenta, TNC is a novel candidate for further study in placental isPTB pathophysiology.

In addition to the upregulation of *C3,* identified by its association with the over-represented IGF signaling pathway, we observed upregulation of other complement related genes, *C1R* and *CFH*. The complement system has a key role in pregnancy, especially during the development of the maternal-fetal interface where the establishment of maternal immunotolerance allows for a semi-allogeneic existence between fetus and mother yet also protect the mother and fetus from infection[34]. Previous studies demonstrated elevated levels of C3a, a cleavage product of C3, at 20 wks gestation was associated with various adverse pregnancy outcomes classified as iatrogenic preterm or sPTB compared to normal term births, however the authors did not distinguish the different subclassification of preterm birth such as preeclampisa (PE), interuterine growth restriction (IUGR), or isPTB[35]. While the upregulation of C3 in our data could be extrinsic to IGF pathway activation, the co-expression and upregulation of C1R, a complement initiator, suggests the activation of the classical pathway [34,35]. However, the corresponding upregulation of CFH, a complement regulator, suggests that an alternative pathway could also be activated, which in the absence of infection could indicate damaged tissue [34,35]. While there is a growing body of evidence, including our data, that associates the complement system with PTB, and especially isPTB, it is unclear if it is a consequence or contributor to the pathology.

In stark contrast to the isPTB signature, the AHC signature we observed was associated with mitotic and cell cycling pathways suggesting active cell division in these placentas. This would be expected at this gestational age range as the placenta is actively remodeling and expanding to accommodate exponential fetal growth[4]. While our AHC candidate gene list from villous tissue does differ from that of Ackerman et al[40], both analyses identified a set of histones upregulated from the *HIST1* locus on Chr6p21-22, suggesting there is active replication occurring in the AHC samples. Furthermore, their pathway analyses included numerous pathways associated with active growth including telomere maintenance, deposition of CENPA and various RNA Pol I related pathways even though their samples are of a much younger gestational age[40]. Together these data do suggest that AHC placentas are in an active growth phase appropriate for their gestational age and that although they are preterm due to infection, they are well suited to be controls for future studies. In contrast, in the isPTB placentas of the same gestational age range as the AHC placentas we did not detect this growth signature, instead there was a lack of significant difference in gene expression between isPTB and TB, suggesting that these placentas are more similar in growth pattern to the term placenta, further supporting advanced placental maturation. Interestingly, the most upregulated gene in the AHC samples, *CDKN1C* (cyclin dependent kinase inhibitor 1C), is located on Chr11p15.5, an essential placental imprinted locus with roles in placental and fetal growth(ref).

The expression pattern observed in our AHC data suggests the imprinting is appropriate in the AHC samples allowing for the transcription of *CDKN1C* from the maternal allele [36]. In contrast to the AHC expression pattern, the PTB and TB patterns are very similar for each gene. Aberrant expression of *Cdkn1c* in mice has been associated with increased fetal and placental growth coupled with changes in placental morphology[37]. Other genes in the locus such as *Phlda2* have been associated with placental overgrowth but without the associated fetal overgrowth in mice[38]. We were not able to assess PHLDA2 expression as it was not well expressed in our data set and subsequently filtered out of the final data set used for analysis. While there is synteny between the mouse and human loci, the mechanisms of regulation via imprinting are different[37,39]. However, given the potential implication on placental specific growth and morphology, it should be considered for further study.

One caveat of the current study, as with other studies, is the lack of normal placentas at early gestational ages which made it necessary to identify appropriate controls. In most studies, gestational aged-matched placentas have an underlying iatrogenic pathology such as PE. Both the Nijman and Morgan studies indicated that the iatrogenic samples commonly used as controls also have MVM and would likely confound the molecular signatures we sought to identify[8,10]. The AHC samples we chose were designated infected by histological examination, but the villi did not appear to be damaged or inflamed, making it an ideal gestational age control for our study. A second caveat is how we identified the infected specimens. The Ackerman et al study confirmed infection through testing of the amniotic fluid. However, our infected specimens were identified by histological analyses. This could explain the differences in our infection related gene lists.

This study is the first to identify transcriptome signatures in placental villous samples from isPTB and gestationally age-matched AHC samples. These signatures highlight the necessity to carefully phenotype sPTB into the respective AHC and isPTB subclasses to pinpoint the potential molecular mechanisms underlying the pathophysiology of isPTB. Ultimately, the insights gained from understanding the differences between AHC and isPTB will improve our ability to differentiate and predict isPTB quite possibly through non-invasive methods. More, importantly, understanding the molecular etiology of isPTB associated with placental hypermaturity may provide translational approaches beyond diagnostics such as treatment of the placenta to ensure appropriate development and birth timing.

## Materials and Methods

### Study Population

This study was approved by the Cincinnati Children’s Hospital Medical Center institutional review board (#IRB 2013-2243, 2015-8030, 2016-2033). De-identified term (n=6), isPTB (n=8), and AHC (n=8) placental villous samples along with appropriate covariate information were obtained from the following sources: The Global Alliance to Prevent Prematurity and Stillbirth (GAPPS) in Seattle Washington USA, the Research Centre for Women’s and Infant’s Health (RCWIH) at Mt Sinai Hospital Toronto Canada, and the University of Cincinnati Medical Center (UCMC). Inclusion criteria included: maternal age 18 years or older, singleton pregnancies with either normal term delivery (38-42 weeks gestation) or preterm delivery (29-36 weeks gestation) without additional complications other than idiopathic spontaneous preterm. Utilizing an RNA sequencing power calculation from[40] (Supplemental Table S1), it was determined that between 5-9 transcriptomes for this particular study to get a powered analysis between 70-90% power at a 2-fold change in expression. We opted for higher power and thus, previously published placental villous transcriptomes generated through RNA sequencing were also utilized (GEO GSE73714 term=5, isPTB=4, inter-amniotic infection=1)[41] bringing our final sample totals to the following: term=11, isPTB=12, and infected=9 (Table 1). Birth weight percentiles were estimated using the WHO weight percentiles calculator with an US average birth weight of 3400gm[42]. It is important to note that the AHC samples were received several months after the study was initiated. Therefore, their transcriptome data were generated after the rest of the data resulting in two batches of data and potential for a batch effect. We have described our assessment and corrections of the batch effects in the respective methods below.

### Clinical definitions

Gestational age was established based on last menstrual period confirmed by an ultrasonographic examination prior to 20 weeks[43]. For samples originating from OSU, infection was established based on analysis of the amniotic fluid retrieved in sterile conditions by trans-abdominal amniocentesis. Amniotic fluid infection (IAI) was established by a positive Gram stain or a positive microbial culture result[44]. For the samples from RCWIH, AHC was established through histologic examination for inflammation of the placenta and fetal membranes which was assessed by a clinical pathologist. Preterm birth was defined as delivery of the neonate <37 weeks GA[45], although we only included samples <36.5 weeks GA. Idiopathic preterm birth was established absent IAI and histologic inflammation of the placenta/fetal membranes as assessed by a clinical pathologist.

### Transcriptome Generation

All placental samples from GAPPS, RCWIH, and UCMC which were collected within 60 minutes of delivery and snap frozen prior to biobanking. Total RNA was prepared from placental villous samples thawed in RNAIce Later (Invitrogen) as per manufacturer instructions. Total RNA was isolated using the RNAeasy Mini Kit (Qiagen). 50-100 μg of total RNA was submitted to the University of Cincinnati Genomics, Epigenomics and Sequencing Core for RNA quality assessment and sequencing. RNA total libraries were generated using a RiboZero Kit (Illumina) and sequencing was run on an Illumina High Seq 2000 system to generate single end 50bp reads at a depth of 50 million reads. Details on the collection of placental samples and generation of transcriptomes from GSE73714 can be found here[41].

### RNA-sequence data preparation

To facilitate RNA sequence analyses, a local instance of the *Galaxy*[46] was utilized with the following tools: *FASTQC* (Galaxy v0.71)[47], *TrimGalore!* (Galaxy v0.4.3.1)[48], *Bowtie2* (Galaxy v2.3.4.1)[49], and *FeatureCounts* (Galaxy v1.6.0.3)[50]. The quality of the raw fastq files was assessed with *FASTQC* and with adapters subsequently trimmed with *TrimGalore*. Trimmed sequences were then aligned to the University of California Santa Cruz (UCSC) human genome hg38 using *Bowtie2.* To avoid cross mapping to paralogous genes expressed in the placental tissues, we set the *Bowtie2* parameters as sensitive end to end. *FeatureCounts* was used to generate total read counts per gene and to generate a count matrix file to be used in differential gene expression analyses.

### Differential Expression Analyses

After annotation, all non-coding transcripts were removed from the count matrix. Using *edgeR* (Emperical Analyses of Digital Gene Expression in R), the count data was then filtered using the counts per million (CPM) method to allow for various quality control analyses to ensure the data was ready for differential expression testing[11]. All data were normalized as a single data set using TMM (trimmed means of m values) to account for differences in library sizes[51]. After normalization, we assessed the data matrix for potential batch effects using principle components analyses (PCA) and did not observe significant batch effects; therefore, no correction was applied to preserve meaningful biological variation. A generalized linear model (glm) was built (using *limma* within the *edgeR* framework) to assess differential gene expression across all birth types accounting for fetal sex. As the entire cohort was small, we did not assess additional covariates for this study. Once the model was generated, we examined the differential expression testing using pairwise contrasts within the model. Multiple corrections testing was performed using the Benjamini Hochberg method with a Q value of <0.05.

### Transcriptomic signature identification

Venny v2.0 was utilized to generate Venn diagrams and identify candidate genes for isPTB and AHC signatures. Candidate genes along with their expression patterns were loaded into Excel and sorted by fold change. The isPTB signature was defined as genes having an upregulated or downregulated expression pattern when compared to AHC or TB and with the AHC vs TB comparison having a non-significant fold change. The AHC signature was defined as genes having upregulated or downregulated expression pattern when compared to isPTB or TB and with the isPTB vs TB comparison having a non-significant fold change. Heatmaps were generated in Prism v8 (GraphPad).

### Immunohistochemistry (IHC)

Immunohistochemistry was performed as previously published[52]. Briefly, all slides were incubated 95°C target retrieval solution for 30 minutes then washed in deionized water. Slides were then incubated in 3% hydrogen peroxide for 10 minutes followed by blocking in 10% normal goat serum +1% bovine albumin for 60 minutes. Primary antibodies generated in rabbit sera were diluted in phosphate buffered saline (PBS): DKK1 (1:20, GeneTex GTX40056) and IGFBP1 (1:50, GeneTex GTX31149). Slides were incubated overnight at 4°C washed and incubated with biotinylated secondary antibody (anti-rabbit) for 60 minutes. Antibody binding was detected using DAB and slides were counterstained with hematoxylin. All slides were imaged on a Nikon Eclipse 80i microscope.

### Signaling Pathway Analyses

Significant genes divided into molecular signature categories were entered into the Panther Pathway DB[53] for statistical overrepresentation analyses for Reactome Pathways. Fisher’s Exact tests were used to determine significance and Bonferroni correction for multiple comparisons. Pathways were considered significant if they had an adjusted p-value <0.05 and enrichment score of >4.

### Statistical Analyses

Data were analyzed in Prism v8 (GraphPad). Data were evaluated for normality and non-parametric tests applied as appropriate. Non-parametric data are expressed as median and range and were analyzed by Kruskal-Wallis Test ANOVA with Dunn’s Multiple Comparisons. Categorical data were analyzed using Fisher’s Exact Test.

## Author Contributions- as per PloS Bio Author Guidelines for authorship

HMB-Conceptualization, project administration, resource acquisition, investigation, formal analyses, data curation, writing of the manuscript including visualization, draft preparation, review and editing.

SGK- Resource acquisition, manuscript review and editing

IAB - Resource acquisition, manuscript review and editing

CSB - Resource acquisitions, manuscript review and editing

WEA- Resource acquisition, manuscript review and editing

LJM^+^- Conceptualization, Funding Acquisition, Resources, Supervision, manuscript review and editing

HNJ^+^- Conceptualization, Resources, Supervision, manuscript draft preparation, review and editing

+ Joint Senior Contributions to the paper.

## Acknowledgements

The authors would like to express their gratitude to the patients who donated their placentas for research and to Pietro Presicce, Paranthaman Senthamarai Kannan, and Manuel Alvarez (Kallapur lab), GAPPS, and RCWIH for assisting us in obtaining the placental samples and covariate data. Additionally, the authors thank the staff at the University of Cincinnati Genomics, Epigenomics and Sequencing Core for their help generating the bulk of the transcriptomics data for this project. This work was supported by grants to LM from the March of Dimes Prematurity Research Center Ohio Collaborative Funding 07/13-6/18, The Bill and Melinda Gates Foundation: Systems Biology Approaches to Birth Timing and Prematurity OPP1113966 Funding 11/14-10/17, and the Eunice Kennedy Shriver National Institute of Child and Health & Human Development of the National Institutes of Health Award Number R01HD091527.

## Data Availability

Transcriptomic data has been made available in the Gene Expression Omnibus (GEO) under accession number: GSE118442 to be released upon publication.

## Additional information

The authors declare they have no competing interests.

