## Supplemental Files for "Unique transcriptomic landscapes identified in idiopathic spontaneous and infection related preterm births compared to normal term births"

**Supplemental Data**

Supplementary Table S1: Power calculations for number of transcriptomes needed for study

| Symbol | Description | 90% power | 80% power | 70% power |
|---|---|---|---|---|
| **Alpha (a)** | Type 1 Error | 0.05 | 0.05 | 0.05 |
| **Beta (b)** | Power | 0.9 | 0.8 | 0.7 |
| **Mu (m)** | Counts[1] | 50 | 50 | 50 |
| **Sigma (s)** | Coefficient of Variation[2] | 0.43 | 0.43 | 0.43 |
| **Delta (D)** | Fold Change (Effect Size) | 2 | 2 | 2 |
| | Number of samples | 9 | 7 | 5 |

[1]Counts equal read depth 50 million is sufficient for this experiment based on previous data
[2]Coefficient of Variation = average human variation calculated from (Hart et al. 2013)

Supplemental Table S2: Candidate genes associated isPTB molecular signature

| | **isPTB vs AHC** | | **isPTB vs TB** | | **AHC vs TB** | |
|---|---|---|---|---|---|---|
| **Gene ID** | **Log2 Fold Change** | **Adjusted P value** | **Log2 Fold Change** | **Adjusted P value** | **Log2 Fold Change** | **Adjusted P value** |
| *PAEP* | 5.3 | 1.32E-03 | 7.1 | 1.50E-05 | 1.8 | 3.83E-01 |
| *IGFBP1* | 5.2 | 8.93E-04 | 7.6 | 3.40E-07 | 2.4 | 1.34E-01 |
| *PRL* | 4.3 | 3.45E-04 | 6.3 | 1.18E-07 | 2.0 | 1.30E-01 |
| *CHRDL1* | 4.0 | 1.24E-02 | 6.8 | 1.11E-05 | 2.8 | 1.06E-01 |
| *GNLY* | 3.9 | 1.87E-05 | 3.9 | 4.67E-06 | -0.01 | 9.97E-01 |
| *GREB1* | 3.9 | 7.78E-05 | 5.1 | 1.18E-07 | 1.2 | 3.22E-01 |
| *RORB* | 3.9 | 5.30E-03 | 6.9 | 3.20E-07 | 3.0 | 4.16E-02 |
| *SCARA5* | 3.8 | 2.32E-04 | 4.3 | 5.48E-06 | 0.6 | 7.03E-01 |
| *CNR1* | 3.7 | 2.86E-04 | 3.3 | 8.76E-04 | -0.4 | 8.27E-01 |

| | | | | | | |
|---|---|---|---|---|---|---|
| *NDP* | 3.4 | 1.47E-02 | 6.4 | 4.52E-06 | 3.0 | 4.76E-02 |
| *HLA-DQA1* | 3.3 | 9.10E-03 | 1.6 | 5.58E-01 | -1.7 | 2.41E-01 |
| *RXFP1* | 3.3 | 1.12E-03 | 4.2 | 1.11E-05 | 0.9 | 4.96E-01 |
| *IGFBP6* | 3.2 | 1.56E-03 | 3.9 | 3.99E-05 | 0.8 | 5.70E-01 |
| *PRUNE2* | 3.1 | 3.48E-04 | 4.0 | 4.80E-07 | 0.9 | 3.84E-01 |
| *DKK1* | 3.0 | 5.49E-06 | 3.8 | 7.16E-10 | 0.8 | 3.68E-01 |
| *MEDAG* | 3.0 | 2.20E-03 | 4.1 | 1.11E-05 | 1.1 | 3.77E-01 |
| *LEFTY2* | 3.0 | 3.03E-03 | 2.4 | 2.31E-02 | -0.5 | 7.38E-01 |
| *RGS1* | 2.9 | 7.78E-05 | 2.1 | 6.87E-03 | -0.8 | 3.72E-01 |
| *RASD1* | 2.8 | 2.88E-04 | 2.8 | 2.08E-04 | -0.03 | 9.85E-01 |
| *TRPC4* | 2.8 | 7.44E-04 | 2.4 | 5.42E-03 | -0.4 | 7.45E-01 |
| *ALDH1A1* | 2.7 | 1.87E-05 | 2.8 | 2.75E-06 | 0.1 | 9.54E-01 |
| *RBP4* | 2.7 | 1.46E-02 | 4.3 | 3.79E-05 | 1.6 | 1.76E-01 |
| *DNASE1L3* | 2.7 | 7.44E-04 | 2.3 | 5.18E-03 | -0.4 | 7.73E-01 |
| *NCAM1* | 2.7 | 3.38E-03 | 2.5 | 7.78E-03 | -0.2 | 9.22E-01 |
| *AHNAK2* | 2.6 | 6.43E-04 | 2.9 | 3.34E-05 | 0.3 | 7.92E-01 |
| *CXCL9* | 2.6 | 9.98E-02 | 3.6 | 2.71E-02 | 1.0 | 6.15E-01 |
| *THBS2* | 2.5 | 3.38E-03 | 2.5 | 3.92E-03 | -0.02 | 9.90E-01 |
| *COL4A4* | 2.5 | 3.08E-03 | 2.8 | 4.48E-04 | 0.4 | 7.72E-01 |
| *CP* | 2.4 | 6.08E-03 | 2.2 | 2.20E-02 | -0.3 | 8.57E-01 |
| *WT1* | 2.4 | 1.27E-02 | 3.5 | 1.42E-04 | 1.2 | 3.21E-01 |
| *WDR72* | 2.4 | 1.54E-02 | 2.5 | 1.47E-02 | 0.2 | 9.32E-01 |
| *RAMP1* | 2.3 | 7.19E-04 | 2.4 | 2.61E-04 | 0.1 | 9.26E-01 |
| *ADAMTS4* | 2.3 | 6.48E-04 | 2.6 | 3.99E-05 | 0.3 | 7.79E-01 |
| *PZP* | 2.3 | 2.43E-03 | 1.8 | 2.88E-02 | -0.5 | 6.70E-01 |
| *TMEM132C* | 2.3 | 5.17E-04 | 2.0 | 2.59E-03 | -0.3 | 8.16E-01 |
| *ALPK2* | 2.2 | 1.43E-05 | 1.3 | 2.05E-02 | -0.9 | 1.78E-01 |
| *ALDH1A2* | 2.2 | 3.29E-04 | 1.9 | 1.69E-03 | -0.3 | 7.99E-01 |
| *TNC* | 2.2 | 3.34E-04 | 2.0 | 1.32E-03 | -0.2 | 8.31E-01 |
| *ABI3BP* | 2.2 | 2.88E-04 | 2.1 | 3.29E-04 | -0.1 | 9.37E-01 |
| *AADAC* | 2.2 | 8.72E-03 | 2.4 | 5.61E-03 | 0.2 | 8.93E-01 |
| *C3* | 2.1 | 2.20E-03 | 2.0 | 5.38E-03 | -0.2 | 8.89E-01 |
| *LSAMP* | 2.1 | 1.75E-02 | 2.5 | 6.83E-03 | 0.4 | 8.05E-01 |
| *GAS1* | 2.1 | 2.05E-04 | 2.7 | 2.58E-07 | 0.6 | 4.33E-01 |
| *CXCL10* | 2.1 | 3.48E-02 | 1.6 | 2.86E-01 | -0.5 | 7.49E-01 |
| *IL1RL1* | 2.1 | 1.08E-02 | 1.1 | 5.14E-01 | -0.9 | 3.50E-01 |
| *RGS22* | 2.1 | 2.56E-03 | 1.2 | 2.91E-01 | -0.9 | 3.05E-01 |
| *CFH* | 2.1 | 2.78E-03 | 1.3 | 1.66E-01 | -0.7 | 4.31E-01 |
| *ITK* | 2.0 | 1.26E-02 | 2.1 | 1.19E-02 | 0.1 | 9.70E-01 |
| *PTGIS* | 2.0 | 1.50E-02 | 1.9 | 4.80E-02 | -0.2 | 9.11E-01 |

| | | | | | | |
|---|---|---|---|---|---|---|
| *CD96* | 2.0 | 3.00E-04 | 1.1 | 1.22E-01 | -0.9 | 2.16E-01 |
| *CILP* | 2.0 | 9.96E-03 | 2.6 | 3.46E-04 | 0.6 | 5.54E-01 |
| *HSD11B1* | 2.0 | 2.95E-03 | 1.6 | 4.02E-02 | -0.4 | 6.78E-01 |
| *MPZL2* | 2.0 | 1.20E-02 | 2.8 | 2.90E-04 | 0.8 | 4.57E-01 |
| *GBP5* | 2.0 | 1.41E-02 | 2.1 | 1.19E-02 | 0.1 | 9.64E-01 |
| *CD69* | 2.0 | 7.19E-04 | 1.2 | 1.51E-01 | -0.8 | 2.67E-01 |
| *OXTR* | 2.0 | 1.55E-03 | 2.3 | 2.20E-04 | 0.3 | 8.04E-01 |
| *KCND2* | 2.0 | 7.84E-02 | 2.8 | 1.37E-02 | 0.8 | 5.73E-01 |
| *CD38* | 2.0 | 4.31E-03 | 2.4 | 3.29E-04 | 0.4 | 6.89E-01 |
| *NAP1L3* | 1.9 | 2.85E-03 | 1.5 | 4.60E-02 | -0.4 | 6.92E-01 |
| *ATOH8* | 1.9 | 6.93E-04 | 1.6 | 6.73E-03 | -0.3 | 7.33E-01 |
| *CYTIP* | 1.9 | 1.32E-03 | 1.0 | 3.78E-01 | -1.0 | 1.92E-01 |
| *ANOS1* | 1.9 | 3.11E-03 | 1.5 | 4.57E-02 | -0.4 | 6.62E-01 |
| *CHI3L2* | 1.9 | 6.37E-04 | 1.6 | 7.78E-03 | -0.3 | 7.14E-01 |
| *GLB1L2* | 1.9 | 2.21E-02 | 2.9 | 2.20E-04 | 1.0 | 3.15E-01 |
| *CXCL11* | 1.9 | 4.54E-02 | 1.3 | 5.08E-01 | -0.6 | 6.34E-01 |
| *PTGS2* | 1.8 | 5.08E-04 | 1.0 | 2.66E-01 | -0.9 | 1.62E-01 |
| *ZBTB16* | 1.8 | 7.00E-03 | 1.2 | 2.57E-01 | -0.6 | 4.78E-01 |
| *OMD* | 1.8 | 2.56E-03 | 2.2 | 1.61E-04 | 0.4 | 7.01E-01 |
| *ISLR* | 1.8 | 6.34E-04 | 2.0 | 5.43E-05 | 0.2 | 8.19E-01 |
| *SMOX* | 1.8 | 3.66E-03 | 2.4 | 3.25E-05 | 0.6 | 4.65E-01 |
| *TNFRSF11B* | 1.8 | 1.73E-02 | 1.3 | 2.27E-01 | -0.4 | 6.96E-01 |
| *TWIST2* | 1.8 | 1.72E-02 | 1.9 | 1.72E-02 | 0.1 | 9.50E-01 |
| *PRRX1* | 1.8 | 1.08E-02 | 1.2 | 2.75E-01 | -0.6 | 5.44E-01 |
| *GEM* | 1.8 | 1.62E-03 | 1.4 | 2.19E-02 | -0.4 | 6.80E-01 |
| *WISP1* | 1.8 | 7.12E-03 | 1.8 | 7.56E-03 | 0.1 | 9.69E-01 |
| *KCNB1* | 1.8 | 6.08E-03 | 1.8 | 5.61E-03 | 0.1 | 9.63E-01 |
| *ADCY1* | 1.7 | 2.11E-02 | 2.3 | 1.69E-03 | 0.6 | 5.76E-01 |
| *SLC12A8* | 1.7 | 2.13E-02 | 2.1 | 5.61E-03 | 0.4 | 7.43E-01 |
| *PRDM1* | 1.7 | 1.63E-03 | 1.9 | 4.84E-04 | 0.1 | 8.92E-01 |
| *OGN* | 1.7 | 3.65E-03 | 1.7 | 5.42E-03 | -0.03 | 9.82E-01 |
| *DEPTOR* | 1.7 | 3.07E-04 | 1.7 | 4.03E-04 | -0.1 | 9.52E-01 |
| *CTSW* | 1.7 | 2.64E-02 | 1.9 | 1.91E-02 | 0.2 | 8.82E-01 |
| *MOV10L1* | 1.7 | 3.85E-02 | 1.1 | 4.87E-01 | -0.6 | 6.04E-01 |
| *EPDR1* | 1.7 | 1.40E-02 | 2.5 | 1.47E-04 | 0.8 | 3.71E-01 |
| *IL7R* | 1.7 | 2.23E-03 | 1.2 | 9.10E-02 | -0.5 | 5.01E-01 |
| *NR4A2* | 1.7 | 7.01E-04 | 1.1 | 6.23E-02 | -0.6 | 4.09E-01 |
| *IGFBP2* | 1.7 | 9.80E-03 | 1.9 | 3.77E-03 | 0.2 | 8.58E-01 |
| *MAATS1* | 1.7 | 8.42E-03 | 1.7 | 1.94E-02 | -0.02 | 9.85E-01 |
| *CLNK* | 1.7 | 1.76E-02 | 1.5 | 6.71E-02 | -0.2 | 8.96E-01 |

| | | | | | | |
|---|---|---|---|---|---|---|
| *PADI1* | 1.7 | 1.93E-01 | 3.1 | 1.12E-02 | 1.4 | 3.36E-01 |
| *ADRA2C* | 1.7 | 1.51E-02 | 1.9 | 7.81E-03 | 0.2 | 8.55E-01 |
| *RBP1* | 1.7 | 6.30E-04 | 1.4 | 4.67E-03 | -0.2 | 7.85E-01 |
| *F2R* | 1.7 | 4.93E-03 | 2.2 | 7.44E-05 | 0.5 | 5.00E-01 |
| *CNTNAP3* | 1.7 | 1.20E-02 | 1.3 | 1.38E-01 | -0.3 | 7.51E-01 |
| *DNER* | 1.6 | 4.24E-02 | 1.3 | 2.80E-01 | -0.3 | 8.11E-01 |
| *MARCO* | 1.6 | 2.90E-02 | 1.0 | 5.88E-01 | -0.7 | 4.87E-01 |
| *TRIM63* | 1.6 | 1.92E-02 | 1.0 | 4.71E-01 | -0.6 | 5.08E-01 |
| *LRRC15* | 1.6 | 1.23E-01 | 2.8 | 7.09E-03 | 1.1 | 3.51E-01 |
| *GGT5* | 1.6 | 1.76E-02 | 1.7 | 1.46E-02 | 0.1 | 9.21E-01 |
| *KLF15* | 1.6 | 1.86E-02 | 1.2 | 2.11E-01 | -0.4 | 7.28E-01 |
| *IL18RAP* | 1.6 | 1.46E-02 | 1.7 | 1.93E-02 | 0.1 | 9.64E-01 |
| *FHOD3* | 1.6 | 2.80E-02 | 2.4 | 3.59E-04 | 0.8 | 3.85E-01 |
| *FNDC4* | 1.6 | 2.93E-03 | 1.7 | 3.92E-03 | 0.1 | 9.64E-01 |
| *IL1R2* | 1.6 | 7.26E-02 | 2.7 | 5.73E-04 | 1.1 | 2.31E-01 |
| *ZNF516* | 1.6 | 1.55E-03 | 1.5 | 2.28E-03 | 0.0 | 9.72E-01 |
| *CPXM1* | 1.6 | 4.09E-03 | 1.6 | 4.67E-03 | 0.005 | 9.96E-01 |
| *SULF1* | 1.6 | 4.39E-03 | 1.2 | 9.48E-02 | -0.4 | 6.10E-01 |
| *C11orf96* | 1.5 | 3.53E-02 | 2.1 | 2.59E-03 | 0.6 | 5.32E-01 |
| *MX2* | 1.5 | 5.08E-04 | 1.1 | 3.76E-02 | -0.5 | 4.53E-01 |
| *SPON2* | 1.5 | 6.60E-03 | 1.6 | 5.18E-03 | 0.1 | 9.32E-01 |
| *KCNQ3* | 1.5 | 3.05E-02 | 1.6 | 3.39E-02 | 0.1 | 9.28E-01 |
| *CD22* | 1.5 | 2.80E-02 | 2.3 | 5.55E-04 | 0.7 | 3.81E-01 |
| *OASL* | 1.5 | 1.08E-02 | 1.2 | 9.67E-02 | -0.3 | 7.65E-01 |
| *PTPN13* | 1.5 | 4.55E-03 | 1.4 | 1.61E-02 | -0.1 | 9.01E-01 |
| *MAOB* | 1.5 | 1.29E-01 | 2.3 | 1.29E-02 | 0.8 | 5.15E-01 |
| *SULF2* | 1.5 | 1.99E-03 | 1.1 | 6.65E-02 | -0.4 | 5.55E-01 |
| *MGST1* | 1.5 | 2.70E-02 | 1.8 | 9.45E-03 | 0.3 | 7.32E-01 |
| *DTNA* | 1.5 | 1.77E-03 | 1.3 | 1.16E-02 | -0.2 | 8.04E-01 |
| *RRAD* | 1.5 | 3.03E-02 | 2.1 | 1.53E-03 | 0.6 | 4.87E-01 |
| *LUM* | 1.4 | 3.54E-03 | 1.5 | 4.67E-03 | 0.004 | 9.97E-01 |
| *WNT9A* | 1.4 | 1.55E-03 | 1.1 | 4.09E-02 | -0.3 | 6.10E-01 |
| *SP140* | 1.4 | 8.90E-03 | 1.0 | 1.96E-01 | -0.4 | 6.11E-01 |
| *SPN* | 1.4 | 4.34E-02 | 1.5 | 5.41E-02 | 0.1 | 9.50E-01 |
| *CCDC141* | 1.4 | 3.67E-02 | 1.5 | 7.83E-02 | 0.047 | 9.73E-01 |
| *RHOH* | 1.4 | 3.69E-02 | 1.3 | 1.34E-01 | -0.1 | 9.01E-01 |
| *COL4A3* | 1.4 | 1.21E-02 | 1.2 | 6.79E-02 | -0.2 | 8.37E-01 |
| *MX1* | 1.4 | 1.50E-03 | 1.1 | 2.25E-02 | -0.3 | 6.84E-01 |
| *IKZF3* | 1.4 | 3.06E-02 | 2.0 | 6.11E-04 | 0.6 | 4.29E-01 |
| *TNFSF10* | 1.4 | 1.92E-02 | 1.2 | 1.24E-01 | -0.2 | 8.32E-01 |

| | | | | | | |
|---|---|---|---|---|---|---|
| *SPSB1* | 1.4 | 1.25E-03 | 1.0 | 3.33E-02 | -0.3 | 5.96E-01 |
| *SPHK1* | 1.4 | 3.20E-02 | 2.0 | 8.49E-04 | 0.7 | 4.15E-01 |
| *ITGAX* | 1.3 | 4.76E-02 | 1.2 | 2.13E-01 | -0.1 | 9.07E-01 |
| *KLHL13* | 1.3 | 1.20E-01 | 1.9 | 3.29E-02 | 0.6 | 5.99E-01 |
| *GPR176* | 1.3 | 2.08E-02 | 1.3 | 6.91E-02 | -0.1 | 9.50E-01 |
| *EGR2* | 1.3 | 4.90E-02 | 1.2 | 1.64E-01 | -0.1 | 9.43E-01 |
| *CHST2* | 1.3 | 1.54E-02 | 2.0 | 4.23E-05 | 0.7 | 2.81E-01 |
| *IGSF10* | 1.3 | 2.59E-02 | 1.5 | 8.35E-03 | 0.2 | 7.91E-01 |
| *HMCN2* | 1.3 | 8.37E-02 | 2.1 | 2.28E-03 | 0.9 | 3.06E-01 |
| *FOXO1* | 1.3 | 9.80E-04 | 1.3 | 6.11E-04 | 0.030 | 9.72E-01 |
| *HAND2* | 1.2 | 3.08E-03 | 1.4 | 8.49E-04 | 0.1 | 8.70E-01 |
| *SLAMF6* | 1.2 | 4.41E-02 | 1.3 | 7.18E-02 | 0.041 | 9.74E-01 |
| *MELTF* | 1.2 | 8.04E-03 | 1.5 | 8.49E-04 | 0.3 | 6.91E-01 |
| *COL8A1* | 1.2 | 7.32E-03 | 1.4 | 2.40E-03 | 0.2 | 8.39E-01 |
| *ALDOC* | 1.2 | 4.73E-02 | 1.3 | 8.75E-02 | 0.1 | 9.55E-01 |
| *GPR132* | 1.2 | 1.25E-02 | 1.0 | 7.17E-02 | -0.2 | 8.54E-01 |
| *CD226* | 1.2 | 1.04E-02 | 1.0 | 6.76E-02 | -0.2 | 8.39E-01 |
| *TIPARP* | 1.2 | 7.63E-04 | 1.1 | 4.19E-03 | -0.1 | 8.42E-01 |
| *GPRIN3* | 1.2 | 1.85E-02 | 1.0 | 9.49E-02 | -0.2 | 8.56E-01 |
| *COL12A1* | 1.2 | 9.92E-02 | 1.6 | 3.32E-02 | 0.4 | 6.59E-01 |
| *PLXNA4* | 1.1 | 3.57E-02 | 1.4 | 1.02E-02 | 0.3 | 7.46E-01 |
| *FKBP5* | 1.1 | 6.55E-02 | 1.4 | 2.17E-02 | 0.3 | 7.06E-01 |
| *CHST7* | 1.1 | 1.62E-02 | 1.2 | 1.01E-02 | 0.1 | 8.72E-01 |
| *CD9* | 1.1 | 1.86E-02 | 1.1 | 2.91E-02 | 0.006 | 9.95E-01 |
| *RASAL3* | 1.1 | 3.85E-02 | 1.0 | 1.51E-01 | -0.1 | 9.16E-01 |
| *CTSK* | 1.1 | 4.16E-03 | 1.0 | 1.97E-02 | -0.1 | 8.57E-01 |
| *MAP3K5* | 1.1 | 4.37E-03 | 1.0 | 1.79E-02 | -0.1 | 8.82E-01 |
| *VEGFA* | 1.1 | 4.40E-02 | 1.3 | 1.41E-02 | 0.3 | 7.48E-01 |
| *KCNJ16* | 1.1 | 2.21E-01 | 1.8 | 4.57E-02 | 0.7 | 4.39E-01 |
| *SHC3* | 1.1 | 2.25E-02 | 1.2 | 1.97E-02 | 0.1 | 8.88E-01 |
| *HAPLN3* | 1.1 | 1.34E-01 | 1.7 | 1.72E-02 | 0.6 | 4.84E-01 |
| *C1R* | 1.1 | 2.80E-02 | 1.2 | 1.72E-02 | 0.1 | 8.73E-01 |
| *ART4* | 1.0 | 2.37E-02 | 1.0 | 8.33E-02 | -0.1 | 9.35E-01 |
| *GALM* | 1.0 | 1.54E-02 | 1.1 | 2.57E-02 | 0.012 | 9.90E-01 |
| *SLC43A3* | 1.0 | 6.70E-02 | 1.2 | 4.74E-02 | 0.2 | 8.15E-01 |
| *CRLF1* | 1.0 | 9.43E-02 | 1.3 | 3.92E-02 | 0.3 | 7.08E-01 |
| *PKDCC* | 1.0 | 4.51E-02 | 1.1 | 6.08E-02 | 0.1 | 9.46E-01 |
| *NFASC* | 1.0 | 1.21E-01 | 1.4 | 4.43E-02 | 0.4 | 6.60E-01 |
| *ISG15* | 1.0 | 1.07E-01 | 1.4 | 2.61E-02 | 0.4 | 6.16E-01 |
| *RUNX3* | 1.0 | 2.64E-01 | 1.7 | 4.40E-02 | 0.7 | 4.43E-01 |

Supplemental Table S3: Candidate genes associated AHC molecular signature

| | **AHC vs TB** | | **AHC vs isPTB** | | **isPTB vs TB** | |
|---|---|---|---|---|---|---|
| **Gene ID** | **Log2 Fold Change** | **Adjusted P value** | **Log2 Fold Change** | **Adjusted P value** | **Log2 Fold Change** | **Adjusted P value** |
| *CDKN1C* | 3.05 | 1.78E-05 | 3.09 | 4.32E-06 | -0.04 | 9.96E-01 |
| *FIBCD1* | 2.49 | 3.67E-09 | 2.78 | 2.39E-12 | -0.29 | 9.26E-01 |
| *CXCL1* | 2.47 | 4.14E-05 | 2.14 | 2.83E-04 | 0.33 | 9.29E-01 |
| *NXPH4* | 2.12 | 7.25E-07 | 2.02 | 7.00E-07 | 0.10 | 9.75E-01 |
| *WNT7B* | 1.85 | 1.07E-04 | 1.35 | 4.67E-03 | 0.50 | 8.49E-01 |
| *BANF1* | 1.80 | 8.36E-12 | 2.01 | 4.26E-16 | -0.21 | 8.95E-01 |
| *RRM2* | 1.66 | 7.87E-07 | 1.36 | 3.76E-05 | 0.31 | 8.83E-01 |
| *CRCT1* | 1.66 | 5.50E-02 | 1.68 | 3.86E-02 | -0.02 | 9.98E-01 |
| *NPIPB4* | 1.62 | 2.82E-02 | 1.43 | 4.07E-02 | 0.32 | 9.15E-01 |
| *REEP6* | 1.62 | 1.45E-02 | 1.35 | 3.55E-02 | 0.27 | 9.52E-01 |
| *MYO1A* | 1.61 | 9.16E-03 | 1.10 | 8.00E-02 | 0.51 | 8.84E-01 |
| *RBPMS2* | 1.57 | 1.69E-03 | 1.17 | 1.96E-02 | 0.39 | 8.88E-01 |
| *H2AFX* | 1.50 | 8.36E-12 | 1.50 | 9.72E-13 | 0.00 | 9.99E-01 |
| *KIF18B* | 1.50 | 6.08E-06 | 1.31 | 4.69E-05 | 0.18 | 9.37E-01 |
| *RPS16* | 1.49 | 1.13E-06 | 1.38 | 3.90E-06 | 0.11 | 9.67E-01 |
| *HMBS* | 1.48 | 2.60E-06 | 1.31 | 1.87E-05 | 0.17 | 9.42E-01 |
| *ZSWIM9* | 1.48 | 2.81E-05 | 1.18 | 6.67E-04 | 0.30 | 8.85E-01 |
| *SAPCD2* | 1.46 | 3.44E-04 | 1.18 | 3.22E-03 | 0.28 | 9.11E-01 |
| *SRM* | 1.44 | 2.38E-06 | 1.16 | 1.05E-04 | 0.28 | 8.77E-01 |
| *CXorf67* | 1.42 | 2.06E-05 | 1.27 | 1.27E-04 | 0.15 | 9.52E-01 |
| *FOXM1* | 1.41 | 4.04E-07 | 1.16 | 1.85E-05 | 0.25 | 8.85E-01 |
| *SUSD2* | 1.41 | 1.04E-05 | 1.62 | 5.55E-08 | -0.21 | 9.13E-01 |
| *FAM83F* | 1.39 | 3.52E-02 | 0.73 | 2.96E-01 | 0.67 | 8.16E-01 |
| *PHOSPHO1* | 1.37 | 2.20E-03 | 1.02 | 1.92E-02 | 0.34 | 8.85E-01 |
| *PPDPF* | 1.36 | 2.38E-03 | 1.00 | 2.75E-02 | 0.36 | 8.85E-01 |
| *HIST1H4A* | 1.34 | 1.98E-04 | 1.44 | 1.87E-05 | -0.10 | 9.72E-01 |
| *DLX4* | 1.34 | 2.44E-05 | 1.15 | 3.08E-04 | 0.19 | 9.27E-01 |
| *PTMS* | 1.32 | 2.81E-06 | 1.12 | 4.06E-05 | 0.19 | 9.09E-01 |
| *CENPA* | 1.31 | 1.04E-03 | 1.01 | 1.04E-02 | 0.30 | 8.90E-01 |
| *TYMS* | 1.30 | 9.28E-08 | 1.33 | 9.88E-09 | -0.02 | 9.91E-01 |
| *PPP1R14A* | 1.30 | 2.97E-02 | 1.18 | 4.07E-02 | 0.12 | 9.75E-01 |
| *CDCA5* | 1.30 | 2.89E-05 | 1.41 | 1.91E-06 | -0.10 | 9.67E-01 |
| *HIST1H2BM* | 1.28 | 5.33E-03 | 0.96 | 3.67E-02 | 0.32 | 8.90E-01 |
| *GTSE1* | 1.28 | 1.57E-05 | 1.27 | 6.98E-06 | 0.00 | 9.99E-01 |
| *TK1* | 1.26 | 6.77E-06 | 1.33 | 5.76E-07 | -0.07 | 9.75E-01 |

| | | | | | | |
|---|---|---|---|---|---|---|
| *CENPM* | 1.24 | 1.97E-03 | 1.40 | 2.39E-04 | -0.15 | 9.63E-01 |
| *HMGA1* | 1.24 | 7.07E-05 | 1.26 | 2.12E-05 | -0.02 | 9.95E-01 |
| *SHCBP1* | 1.23 | 4.82E-05 | 1.23 | 1.87E-05 | -0.01 | 9.99E-01 |
| *CDC20* | 1.22 | 9.18E-05 | 1.10 | 3.28E-04 | 0.12 | 9.63E-01 |
| *CDT1* | 1.22 | 4.75E-06 | 1.27 | 5.62E-07 | -0.06 | 9.78E-01 |
| *E2F7* | 1.21 | 1.01E-04 | 1.01 | 9.26E-04 | 0.20 | 9.24E-01 |
| *TICRR* | 1.20 | 1.16E-05 | 1.12 | 2.07E-05 | 0.08 | 9.74E-01 |
| *ESPL1* | 1.20 | 1.69E-04 | 0.86 | 8.04E-03 | 0.34 | 8.38E-01 |
| *UHRF1* | 1.19 | 3.52E-03 | 0.90 | 2.72E-02 | 0.30 | 8.90E-01 |
| *IGF2* | 1.19 | 5.09E-03 | 0.79 | 7.52E-02 | 0.40 | 8.59E-01 |
| *MAGEA4* | 1.18 | 1.05E-01 | 1.85 | 7.47E-03 | -0.67 | 8.49E-01 |
| *FAM111B* | 1.18 | 2.71E-02 | 0.86 | 1.02E-01 | 0.32 | 9.11E-01 |
| *RNF26* | 1.17 | 3.00E-09 | 1.24 | 3.58E-11 | -0.06 | 9.71E-01 |
| *TICAM1* | 1.17 | 8.92E-06 | 1.00 | 1.29E-04 | 0.17 | 9.21E-01 |
| *CDKN2A* | 1.17 | 2.16E-03 | 0.88 | 2.14E-02 | 0.29 | 8.90E-01 |
| *QDPR* | 1.17 | 8.91E-04 | 0.93 | 8.19E-03 | 0.24 | 9.00E-01 |
| *PCLAF* | 1.17 | 2.00E-02 | 1.37 | 2.68E-03 | -0.20 | 9.52E-01 |
| *PSAT1* | 1.17 | 1.16E-02 | 1.10 | 1.28E-02 | 0.07 | 9.85E-01 |
| *MYBL2* | 1.15 | 4.48E-05 | 1.19 | 1.32E-05 | -0.03 | 9.88E-01 |
| *PLK1* | 1.15 | 1.48E-06 | 1.11 | 1.70E-06 | 0.04 | 9.81E-01 |
| *CDC25A* | 1.15 | 2.26E-04 | 1.28 | 1.55E-05 | -0.13 | 9.55E-01 |
| *HIST1H3G* | 1.15 | 5.65E-04 | 1.07 | 8.03E-04 | 0.08 | 9.75E-01 |
| *PRR11* | 1.13 | 1.02E-04 | 1.02 | 3.28E-04 | 0.11 | 9.63E-01 |
| *FAM83D* | 1.13 | 2.67E-04 | 1.07 | 3.40E-04 | 0.06 | 9.80E-01 |
| *MKI67* | 1.13 | 9.23E-05 | 1.00 | 4.23E-04 | 0.13 | 9.50E-01 |
| *HRCT1* | 1.13 | 3.96E-02 | 0.98 | 6.19E-02 | 0.15 | 9.72E-01 |
| *HIST1H3H* | 1.13 | 7.89E-04 | 0.89 | 7.56E-03 | 0.23 | 8.94E-01 |
| *BIRC5* | 1.12 | 2.05E-03 | 1.24 | 2.95E-04 | -0.11 | 9.67E-01 |
| *FJX1* | 1.12 | 8.14E-03 | 0.98 | 1.79E-02 | 0.15 | 9.63E-01 |
| *HIST1H3I* | 1.12 | 1.36E-03 | 0.90 | 1.00E-02 | 0.23 | 9.11E-01 |
| *KLF16* | 1.12 | 1.16E-05 | 0.89 | 5.00E-04 | 0.23 | 8.84E-01 |
| *WNK2* | 1.12 | 1.12E-03 | 0.68 | 6.01E-02 | 0.44 | 6.95E-01 |
| *ATN1* | 1.12 | 4.94E-04 | 0.93 | 3.60E-03 | 0.19 | 9.27E-01 |
| *SLC25A29* | 1.12 | 1.94E-03 | 0.75 | 4.36E-02 | 0.37 | 8.36E-01 |
| *CNOT3* | 1.11 | 4.87E-02 | 0.92 | 8.65E-02 | 0.19 | 9.63E-01 |
| *FOXI3* | 1.11 | 9.74E-04 | 1.07 | 8.41E-04 | 0.04 | 9.88E-01 |
| *HIST1H2AL* | 1.10 | 1.86E-03 | 1.06 | 1.67E-03 | 0.04 | 9.88E-01 |
| *CDH3* | 1.10 | 2.26E-03 | 1.01 | 3.97E-03 | 0.09 | 9.75E-01 |
| *SEPHS2* | 1.10 | 5.34E-08 | 1.13 | 5.39E-09 | -0.03 | 9.88E-01 |
| *SGO1* | 1.10 | 4.78E-03 | 0.92 | 1.54E-02 | 0.18 | 9.47E-01 |

| | | | | | | |
|---|---|---|---|---|---|---|
| *ILDR1* | 1.10 | 6.79E-03 | 1.06 | 5.23E-03 | 0.03 | 9.93E-01 |
| *FAM167B* | 1.09 | 4.21E-02 | 1.07 | 2.91E-02 | 0.02 | 9.98E-01 |
| *AURKB* | 1.08 | 1.16E-03 | 1.11 | 4.65E-04 | -0.03 | 9.93E-01 |
| *PMVK* | 1.08 | 2.96E-04 | 1.18 | 2.23E-05 | -0.10 | 9.67E-01 |
| *BICRA* | 1.08 | 4.15E-04 | 0.87 | 4.78E-03 | 0.21 | 8.97E-01 |
| *TPX2* | 1.08 | 1.83E-04 | 1.05 | 1.61E-04 | 0.03 | 9.88E-01 |
| *CCNF* | 1.08 | 1.45E-05 | 0.87 | 4.21E-04 | 0.20 | 8.88E-01 |
| *RAD51* | 1.07 | 4.75E-06 | 1.22 | 3.28E-08 | -0.14 | 9.27E-01 |
| *SP6* | 1.07 | 3.18E-04 | 1.21 | 1.68E-05 | -0.14 | 9.48E-01 |
| *KIF4A* | 1.07 | 1.87E-03 | 0.90 | 7.12E-03 | 0.16 | 9.47E-01 |
| *TP53I11* | 1.06 | 8.93E-04 | 0.64 | 5.92E-02 | 0.43 | 6.67E-01 |
| *PKMYT1* | 1.06 | 1.32E-03 | 1.01 | 1.48E-03 | 0.05 | 9.85E-01 |
| *GET4* | 1.06 | 3.10E-02 | 0.78 | 1.13E-01 | 0.28 | 9.16E-01 |
| *CLIP3* | 1.06 | 2.38E-02 | 0.79 | 8.82E-02 | 0.27 | 9.18E-01 |
| *LRFN3* | 1.06 | 1.87E-02 | 0.63 | 1.81E-01 | 0.43 | 8.49E-01 |
| *CDCP1* | 1.05 | 2.97E-02 | 0.67 | 1.80E-01 | 0.38 | 8.75E-01 |
| *CLSPN* | 1.05 | 3.59E-03 | 1.11 | 8.68E-04 | -0.06 | 9.80E-01 |
| *RAB11B* | 1.04 | 5.84E-06 | 1.03 | 4.45E-06 | 0.01 | 9.95E-01 |
| *KRT10* | 1.04 | 5.15E-04 | 0.90 | 2.39E-03 | 0.13 | 9.51E-01 |
| *CMBL* | 1.04 | 5.33E-03 | 0.88 | 1.43E-02 | 0.15 | 9.52E-01 |
| *ADAMTS19* | 1.04 | 6.21E-03 | 1.03 | 3.42E-03 | 0.01 | 9.98E-01 |
| *HIST1H1B* | 1.04 | 1.12E-03 | 0.93 | 2.62E-03 | 0.11 | 9.63E-01 |
| *CDCA2* | 1.03 | 4.58E-03 | 1.11 | 1.12E-03 | -0.08 | 9.77E-01 |
| *RTL8A* | 1.03 | 1.77E-04 | 1.07 | 4.69E-05 | -0.04 | 9.86E-01 |
| *SLC1A2* | 1.03 | 3.17E-02 | 0.66 | 1.74E-01 | 0.37 | 8.80E-01 |
| *HPRT1* | 1.02 | 2.45E-04 | 0.94 | 5.52E-04 | 0.08 | 9.74E-01 |
| *MT2A* | 1.02 | 2.20E-03 | 0.71 | 3.88E-02 | 0.31 | 8.60E-01 |
| *SOX14* | 1.02 | 2.98E-03 | 1.00 | 2.25E-03 | 0.02 | 9.96E-01 |
| *ZNF865* | 1.02 | 1.12E-03 | 0.84 | 6.56E-03 | 0.17 | 9.29E-01 |
| *FOXP4* | 1.01 | 5.09E-05 | 0.70 | 6.64E-03 | 0.31 | 7.54E-01 |
| *CDC45* | 1.01 | 6.12E-03 | 0.87 | 1.51E-02 | 0.14 | 9.59E-01 |
| *LMNB1* | 1.01 | 8.51E-04 | 0.93 | 1.39E-03 | 0.08 | 9.75E-01 |
| *CCDC86* | 1.01 | 3.78E-03 | 0.80 | 2.03E-02 | 0.21 | 9.25E-01 |
| *CKAP2L* | 1.00 | 5.15E-04 | 0.91 | 1.12E-03 | 0.09 | 9.67E-01 |
| *ZNF703* | 1.00 | 1.26E-02 | 1.20 | 1.18E-03 | -0.19 | 9.36E-01 |
| *IGDCC3* | 1.00 | 8.91E-04 | 1.08 | 1.29E-04 | -0.08 | 9.72E-01 |
| *POLR3K* | 0.98 | 7.83E-04 | 1.02 | 2.30E-04 | -0.04 | 9.86E-01 |
| *CCNA2* | 0.97 | 5.44E-04 | 1.14 | 1.32E-05 | -0.17 | 9.16E-01 |
| *UBE2C* | 0.97 | 2.78E-03 | 1.24 | 2.61E-05 | -0.27 | 8.84E-01 |
| *ASF1B* | 0.96 | 2.16E-03 | 1.08 | 2.81E-04 | -0.11 | 9.63E-01 |

| | | | | | | |
|---|---|---|---|---|---|---|
| *LYPD3* | 0.96 | 1.26E-02 | 1.08 | 2.56E-03 | -0.12 | 9.67E-01 |
| *SKA3* | 0.94 | 4.38E-03 | 1.11 | 2.93E-04 | -0.17 | 9.33E-01 |
| *ANP32B* | 0.93 | 1.32E-06 | 1.05 | 5.39E-09 | -0.12 | 9.26E-01 |
| *PTTG1* | 0.92 | 6.53E-03 | 1.10 | 4.00E-04 | -0.18 | 9.29E-01 |
| *CENPW* | 0.91 | 3.31E-02 | 1.02 | 9.14E-03 | -0.11 | 9.74E-01 |
| *AP1S3* | 0.89 | 7.58E-03 | 1.03 | 7.83E-04 | -0.14 | 9.51E-01 |
| *PBK* | 0.88 | 3.10E-02 | 1.03 | 5.25E-03 | -0.16 | 9.52E-01 |
| *NETO2* | 0.87 | 3.01E-02 | 1.00 | 5.87E-03 | -0.13 | 9.63E-01 |
| *SMAGP* | 0.87 | 2.05E-03 | 1.04 | 7.46E-05 | -0.16 | 9.25E-01 |
| *AP3B2* | 0.86 | 1.43E-01 | 1.30 | 1.33E-02 | -0.43 | 8.84E-01 |
| *LARGE2* | 0.83 | 1.83E-02 | 1.09 | 4.62E-04 | -0.27 | 8.85E-01 |
| *RPS21* | 0.83 | 1.30E-03 | 1.02 | 1.87E-05 | -0.19 | 8.90E-01 |
| *FZD10* | 0.80 | 1.25E-01 | 1.05 | 2.19E-02 | -0.25 | 9.27E-01 |
| *CDCA8* | 0.80 | 7.96E-03 | 1.07 | 9.58E-05 | -0.27 | 8.61E-01 |
| *DUSP9* | 0.77 | 8.79E-02 | 1.04 | 7.74E-03 | -0.27 | 8.95E-01 |
| *FBXO24* | 0.74 | 1.78E-01 | 1.03 | 3.14E-02 | -0.30 | 9.21E-01 |
| *BCAM* | 0.71 | 1.54E-02 | 1.02 | 1.11E-04 | -0.30 | 8.04E-01 |
| *LAMA1* | 0.69 | 1.67E-01 | 1.12 | 7.12E-03 | -0.43 | 8.42E-01 |
| *CPS1* | 0.67 | 1.94E-01 | 1.04 | 1.65E-02 | -0.37 | 8.68E-01 |
| *CLCA2* | 0.66 | 2.76E-01 | 1.02 | 4.70E-02 | -0.36 | 8.90E-01 |
| *KRTAP26-1* | 0.50 | 4.49E-01 | 1.13 | 3.59E-02 | -0.62 | 7.14E-01 |
| *HTR1D* | 0.49 | 3.89E-01 | 1.02 | 2.52E-02 | -0.53 | 7.55E-01 |
| *CFAP54* | -0.96 | 1.02E-01 | -1.32 | 1.01E-02 | 0.36 | 8.90E-01 |
| *PKD1L1* | -0.97 | 5.63E-02 | -1.03 | 2.97E-02 | 0.06 | 9.87E-01 |
| *MPP3* | -0.98 | 6.45E-02 | -1.07 | 2.73E-02 | 0.10 | 9.76E-01 |
| *MEGF10* | -1.01 | 3.67E-02 | -1.15 | 9.55E-03 | 0.14 | 9.67E-01 |
| *DAPP1* | -1.02 | 1.92E-02 | -0.87 | 4.18E-02 | -0.14 | 9.63E-01 |
| *FGL1* | -1.03 | 4.67E-02 | -1.00 | 3.93E-02 | -0.02 | 9.96E-01 |
| *SSBP1* | -1.03 | 7.63E-04 | -0.70 | 2.75E-02 | -0.33 | 8.00E-01 |
| *AMY2B* | -1.03 | 1.53E-03 | -0.97 | 2.24E-03 | -0.06 | 9.79E-01 |
| *GBGT1* | -1.04 | 4.66E-03 | -0.84 | 2.11E-02 | -0.20 | 9.26E-01 |
| *CFAP58* | -1.04 | 1.13E-02 | -0.90 | 2.90E-02 | -0.14 | 9.62E-01 |
| *PILRA* | -1.05 | 6.84E-02 | -1.47 | 4.06E-03 | 0.42 | 8.77E-01 |
| *MYBPHL* | -1.07 | 4.30E-02 | -0.90 | 9.10E-02 | -0.17 | 9.63E-01 |
| *LAMB4* | -1.08 | 1.18E-02 | -0.96 | 2.40E-02 | -0.12 | 9.67E-01 |
| *C11orf52* | -1.08 | 3.97E-02 | -0.77 | 1.55E-01 | -0.31 | 9.00E-01 |
| *C2CD6* | -1.09 | 4.33E-02 | -0.93 | 8.07E-02 | -0.16 | 9.63E-01 |
| *GALNT15* | -1.12 | 1.05E-01 | -1.52 | 1.27E-02 | 0.40 | 8.94E-01 |
| *CASQ1* | -1.12 | 2.64E-01 | -2.00 | 2.18E-02 | 0.88 | 7.92E-01 |
| *XCL1* | -1.14 | 8.41E-03 | -0.99 | 2.19E-02 | -0.15 | 9.58E-01 |

| | | | | | | |
|---|---|---|---|---|---|---|
| *FCN1* | -1.17 | 2.35E-02 | -0.89 | 8.10E-02 | -0.27 | 9.21E-01 |
| *WDR49* | -1.17 | 8.46E-03 | -1.05 | 1.69E-02 | -0.12 | 9.67E-01 |
| *DDI1* | -1.30 | 6.46E-02 | -1.69 | 7.89E-03 | 0.39 | 9.10E-01 |
| *ANG* | -1.41 | 2.97E-02 | -1.47 | 1.56E-02 | 0.06 | 9.88E-01 |
| *ZNF593* | -1.45 | 1.61E-02 | -1.30 | 2.72E-02 | -0.15 | 9.71E-01 |
| *ABCA9* | -1.51 | 1.22E-04 | -1.59 | 2.52E-05 | 0.09 | 9.75E-01 |
| *HBA2* | -1.56 | 2.96E-02 | -1.57 | 1.73E-02 | 0.01 | 9.99E-01 |
| *GAPT* | -1.57 | 2.05E-02 | -1.43 | 2.99E-02 | -0.14 | 9.75E-01 |
| *PPP1R2B* | -1.59 | 1.45E-05 | -1.39 | 2.03E-04 | -0.20 | 9.27E-01 |
| *ABCA6* | -1.62 | 1.03E-04 | -1.86 | 3.94E-06 | 0.24 | 9.26E-01 |
| *HBB* | -1.80 | 3.99E-03 | -1.56 | 9.14E-03 | -0.24 | 9.52E-01 |
| *ADH1B* | -1.93 | 5.82E-03 | -1.73 | 1.09E-02 | -0.20 | 9.67E-01 |
| *ACSM5* | -2.04 | 5.15E-04 | -2.45 | 6.26E-06 | 0.41 | 8.85E-01 |
| *MAP1LC3C* | -2.25 | 1.79E-04 | -2.67 | 5.03E-06 | 0.42 | 8.90E-01 |
| *ADAMDEC1* | -2.94 | 1.97E-04 | -3.31 | 1.68E-05 | 0.36 | 9.39E-01 |
